# Generation of human iPSC-derived pancreatic organoids to study pancreas development and disease

**DOI:** 10.1101/2024.10.24.620102

**Authors:** Jean-Francois Darrigrand, Abigail Isaacson, Francesca M. Spagnoli

## Abstract

The pancreas has vital endocrine and exocrine functions that can be affected by life-threatening diseases such as diabetes and pancreatic cancer. While animal models have been essential for understanding pancreatic development and disease, they are limited by their low-throughput and major species-specific molecular and physiological differences. Generating 3D in vitro models, such as organoids, that are physiologically relevant is thus primordial for investigating pancreatic development and disease in the human context. However, producing human stem cell-derived pancreatic organoids with proper branched architecture and correct patterning of cell domains has remained challenging. Here, we successfully developed a protocol that efficiently and reproducibly generates such organoids from human induced pluripotent stem cells (hiPSCs) by optimising organoid culture format and media. Our differentiation protocol promotes acinar cell differentiation and generates organoids with branches patterned into central trunk and peripheral tip domains, without relying on animal-derived matrices for organoid culture. This platform opens the door to high-throughput investigations of human pancreas development in a system recapitulating the most important aspects of pancreatic tissue architecture. Lastly, we anticipate that this system will contribute to the partial replacement of animal models used to investigate diseases such as pancreatic cancer.

## Introduction

The pancreas has two vital functions, endocrine and exocrine, which control digestion and blood glucose levels. The exocrine compartment makes up over 90% of the organ and acinar cells are its primary constituent^1^. They are responsible for producing digestive enzymes, such as amylase, protease and lipase, stored as zymogen granules and released by exocytosis during digestion. Also, as part of the exocrine compartment, ductal cells form a ramified network of branches that transport the digestive enzymes to the duodenum and release bicarbonate ions to neutralize the acidic contents of the stomach.

During embryogenesis, the pancreas originates from the foregut endoderm at embryonic day (E) 9.5 in mice and 30 days post-conception (dpc) in humans^2,3^. Until E14.5 in mice and 50dpc in human, primary branches form and elongate^4,5^. After E14.5, branches start bifurcating and form a ramified network of ducts connecting all acini to the duodenum. Concomitant with branch formation, pancreatic cells acquire specialized identities and localize into discrete regions of the developing pancreas. Bipotent progenitor cells located in the trunk regions give rise to ductal and endocrine cells, while progenitors in the tip regions give rise to acinar cells^6,7^. Hence, the cellular mechanisms regulating cell differentiation and branching morphogenesis are spatiotemporally synchronized so that pancreatic cells organize themselves into a branched architecture fundamental for pancreatic function.

The gene regulatory networks (GRNs) governing pancreatic identities have been extensively characterized in mice and appear conserved in humans to some extent^8–10^. The transcription factors *PTF1A* and *NKX6*.*1* respectively govern the allocation of multipotent progenitors to either an acinar fate or ductal/endocrine cell identities^9^. However, the role of many soluble and extracellular matrix proteins present in the pancreatic microenvironment remain elusive. The extended time required to study the effects of gene mutations in mice has impeded the systematic characterization of developmental signals. Moreover, clear differences have now been described between human and mouse pancreas development highlighting the importance of developing human experimental models^11^, which would also enable high-throughput functional characterizations.

The *in vitro* differentiation of human induced pluripotent stem cells (hiPSCs) followed by cultures in three-dimension (3D) can generate cellular structures, called organoids, which recapitulate key physiological organ features^12,13^. The use of hiPSCs-derived organoids enables the systematic targeting of developmental pathways by using CrispR-Cas9 technology^14^. Most of the organoid culture protocols rely however on animal-derived materials, such as Matrigel, which are used as scaffolds to support organoid growth and morphogenesis. Here, we optimised a Matrigel-free 3D culture approach generating hiPSC-derived pancreas organoids with a branched architecture and proper regionalisation of acinar/tip and trunk compartments.

Understanding what regulates acinar cell differentiation is essential not only for investigating the molecular mechanisms underlying pancreas development but also those underlying pancreatic cell plasticity in disease. Most importantly, acinar cell plasticity is linked to pancreatic inflammation and pancreatic ductal adenocarcinoma (PDAC). During pancreatic inflammation, acinar cells might dedifferentiate into ductal-like progenitor cells, which reacquire clear embryonic markers^15^. This is a reversible process that allows the regeneration of pancreatic tissue after inflammation. However, persistent activation of mutated genes, such as oncogenic *Kras*, leads to permanent dedifferentiation of acinar cells and is linked to PDAC onset in 98% of cases^1^. The animal procedures used to model pancreatic inflammation and PDAC range from moderate to severe^16,17^. Beside partially replacing the use of such animal procedures, human pancreatic organoids hold great promise for drug screening purposes. The organoid models developed previously either lacked a physiologically relevant architecture or still required to be transplanted in mice and thus remained low-throughput and dependent on animal surgical procedures^18^. Hence, we anticipate that the physiological relevance of the pancreatic organoid platform that we developed here could support the development of high-throughput preclinical studies while partially replacing mouse models of pancreatic diseases.

## Methods

### hiPSC culture

hiPSCs (AICS-0090-39)^19^ were maintained according to the manufacturer’s protocol (https://www.coriell.org/0/PDF/Allen/ipsc/AICS-0090-391_CofA.pdf) on Matrigel-coated (Corning) 6 well-plates with mTesr1 growth medium (STEMCELL Technologies) supplemented with 1% Penicillin-Streptomycin (P/S) in a humidified incubator (37 °C, 5 % CO2). hiPSCs were thawed at 37°C in a water bath and the cell suspension was then transferred dropwise to a falcon tube containing enough growth medium for a 1:10 dilution. hiPSCs were centrifuged, resuspended in growth medium containing 5μM ROCK inhibitor, and then seeded onto Matrigel-coated plates for maintenance.

Cells were passaged as single-cells at 70-80% confluency using Accutase (Invitrogen) and medium supplemented with 5μM Rho-associated protein kinase (ROCK) inhibitor Y-27632 (Sigma) on the day of passaging. Cryopreservation of hiPSCs was carried out in freezing medium comprising 60% mTesr1, 30% knockout serum replacement (KoSR) and 10% dimethyl sulphoxide for 24h at -80°C using controlled-rate freezing. Vials of cells were transferred to liquid nitrogen for long-term storage.

### hiPSC differentiation into pancreatic progenitors

hiPSCs were seeded at a density of 200,000-300,000 cells/cm2 on Matrigel-coated 6-well plates and grown for 24h in mTesr1 supplemented with 1% P/S and 5 μM ROCK inhibitor. Cells were subsequently differentiated in medium that was replenished daily and supplemented with cytokines to induce their specification into pancreatic progenitors. Cytokine and small compound concentrations used were as described in Barsby et al^20^.

Briefly, the first basal medium (Basal Medium 1) used between day 1 and 5 of differentiation was made of MCDB131 medium supplemented with Glutamax (1x), Glucose (2.5 M), NaHCO3 (1.5g/l) and BSA (0.5%). The second basal medium (Basal Medium 2) used between day 6 to the end of differentiation was made of MCDB131 medium supplemented with Glutamax (1x), Glucose (2.5 M), NaHCO3 (2.5g/l), BSA (2%) and ITS-X (0.5x). The reagents required are listed in Table 1.

**Table 1.** List of reagents and suppliers.

| Reagents | Source |
| --- | --- |
| MCDB131 Medium | Gibco (Cat # 10372-019) |
| Glutamax | Gibco (Cat # 35050-038) |
| Glucose | Roth (Cat # X997.1) |
| NaHCO<sub>3</sub> | Gibco (Cat # 11360-039) |
| BSA | Sigma (Cat # A1470) |
| ITS-X | Gibco (Cat # 41400-045) |
| Pen/Strep | Gibco (Cat # 15140122) |
| Activin A (AA) | Qkine (Cat # QK001) |
| CHIR-99021 (Chir) | Abcam (Cat # ab120890) |
| FGF7 | Qkine (Cat # QK046) |
| Vitamin C (Vit. C) | Merck (Cat # A4544) |
| Sant1 | Merck (Cat # S4572) |
| Retinoic Acid (RA) | Sigma (Cat # R2625) |
| LDN-193189 (LDN) | Stemgent (Cat # 04-0074) |
| TPB | Calbiochem (Cat # 565740) |
| EGF | Qkine (Cat # QK011) |
| Nicotinamide (Nicot.) | Sigma (Cat # N0636) |
| ROCK inhibitor Y-27632 (RI) | Sigma (Cat # Y0503) |
| GSiXX | Millipore (Cat # 565790) |
| FGF10 | Qkine (Cat # QK003) |

The supplements used to differentiate pancreatic progenitors were iteratively added to the basal media as detailed in the Table 2.

**Table 2.**
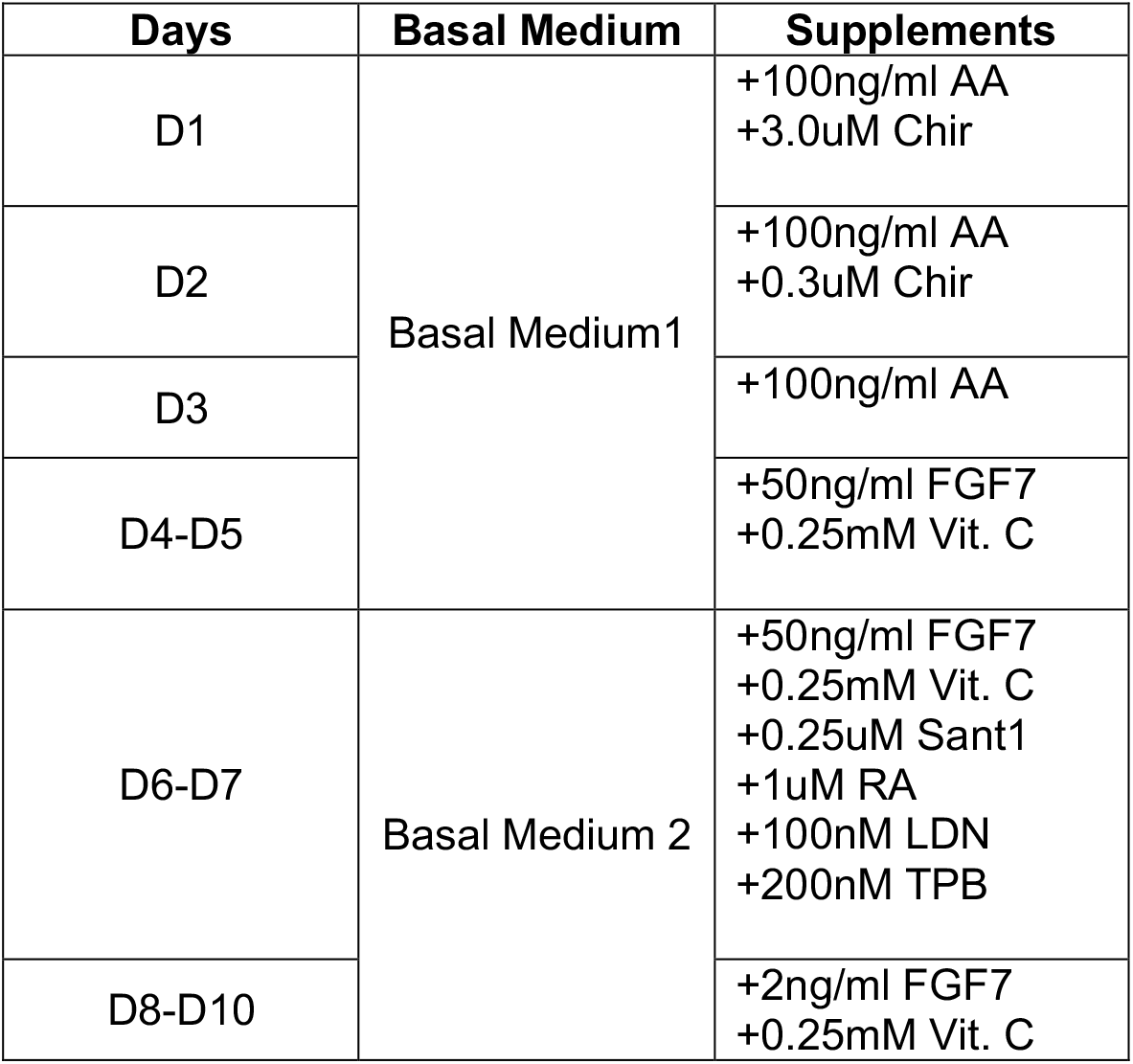
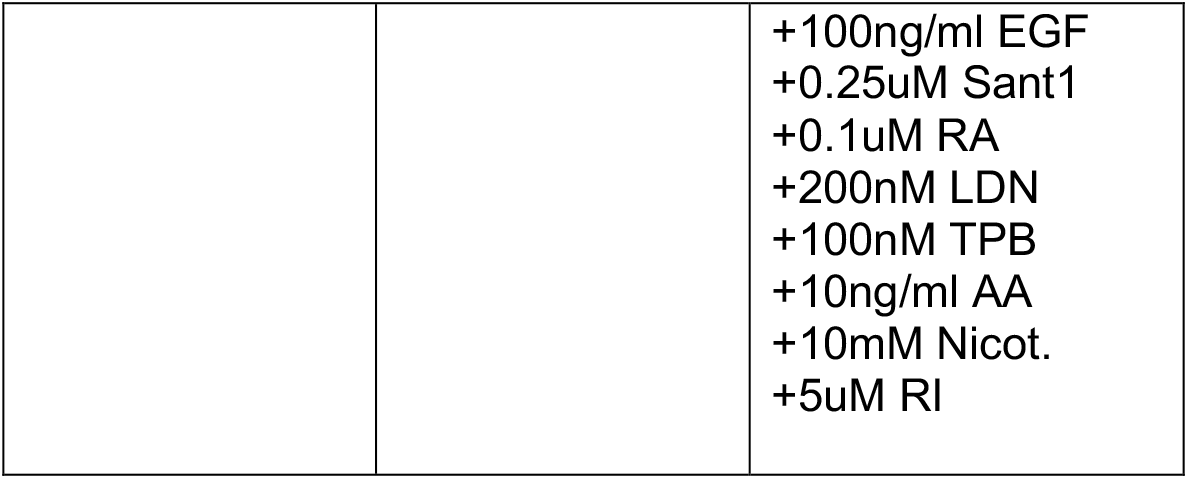
Overview of the supplements and concentrations used to differentiate hiPSCs into pancreatic progenitors.

| Days | Basal Medium | Supplements |
| --- | --- | --- |
| D1 | Basal Medium1 | +100ng/ml AA<br>+3.0uM Chir |
| D2 |  | +100ng/ml AA<br>+0.3uM Chir |
| D3 |  | +100ng/ml AA |
| D4-D5 |  | +50ng/ml FGF7<br>+0.25mM Vit. C |
| D6-D7 | Basal Medium 2 | +50ng/ml FGF7<br>+0.25mM Vit. C<br>+0.25uM Sant1<br>+1uM RA<br>+100nM LDN<br>+200nM TPB |
| D8-D10 |  | +2ng/ml FGF7<br>+0.25mM Vit. C |
|  |  | +100ng/ml EGF<br>+0.25uM Sant1<br>+0.1uM RA<br>+200nM LDN<br>+100nM TPB<br>+10ng/ml AA<br>+10mM Nicot.<br>+5uM RI |

### Formation and culture of pancreatic organoids

On Day 11 of differentiation pancreatic progenitors (PP) were dissociated using TrypLE (Gibco) and left to aggregate in the Aggrewell microwell system (STEMCELL Technologies) for 72h. Progenitors were seeded in the Aggrewell at a density of 1200 cells/microwell. After PP clusters had formed in the Aggrewell (72h), these were transferred to suspension plates, and the basal medium was supplemented with the cytokines and small compounds listed in Table 3.

**Table 3.**
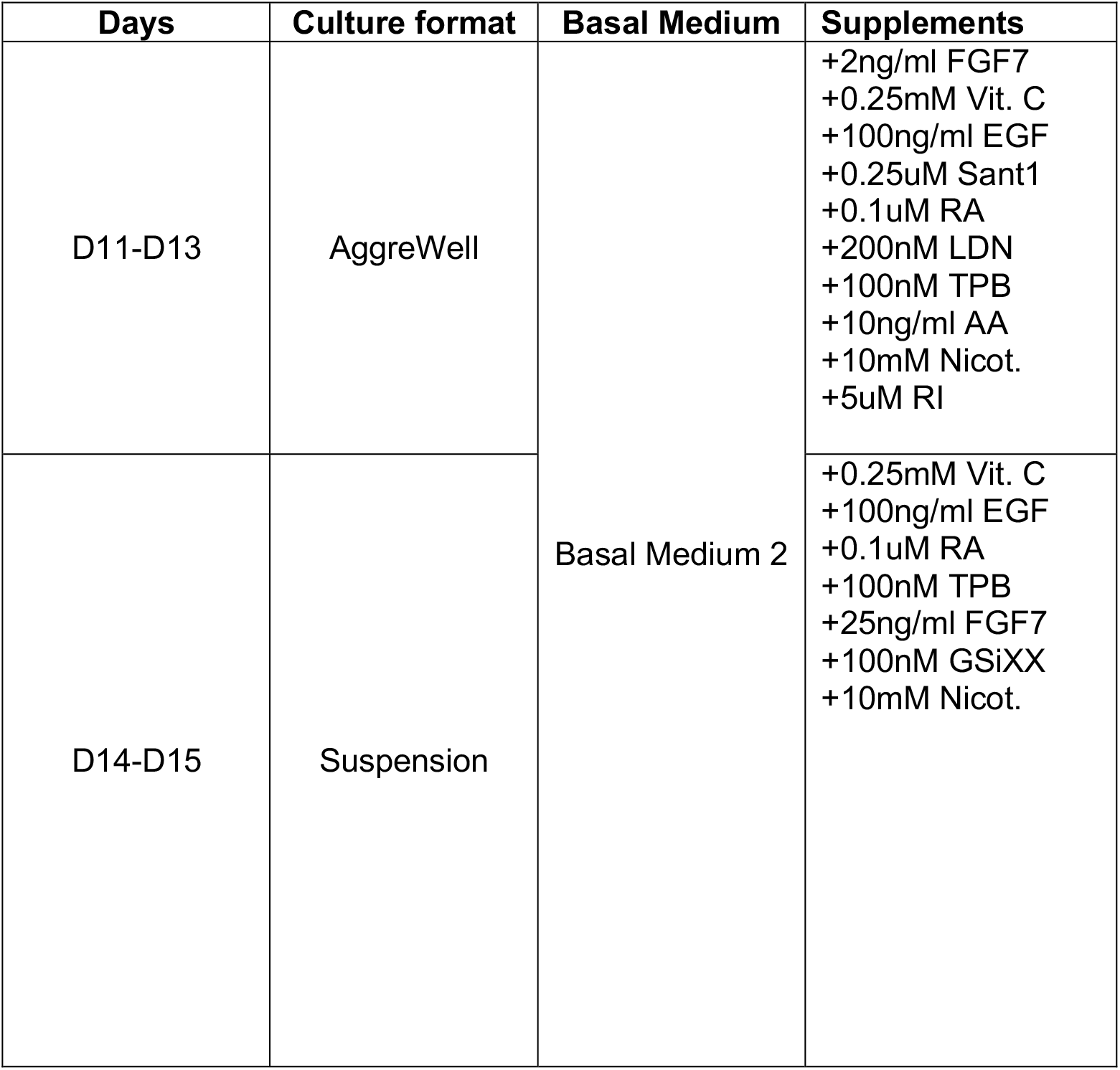
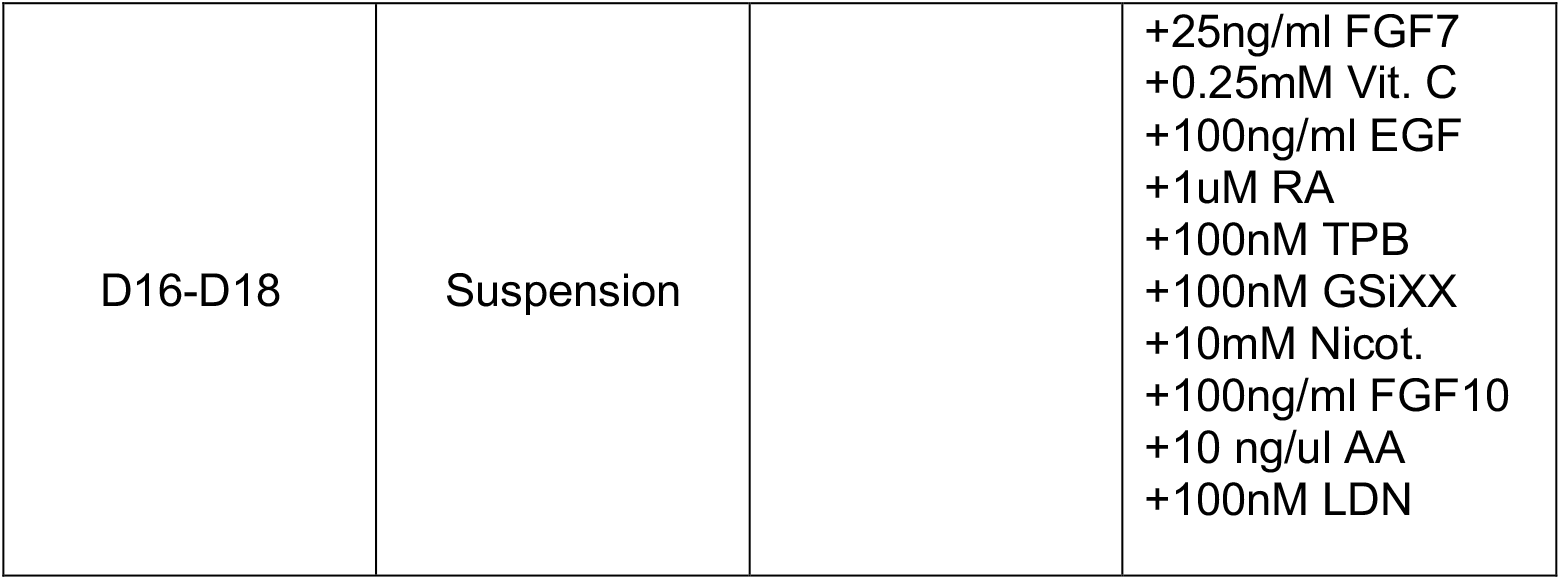
Overview of the supplements and concentrations used to differentiate pancreatic progenitors into branched pancreatic organoids.

| Days | Culture format | Basal Medium | Supplements |
| --- | --- | --- | --- |
| D11-D13 | AggreWell | Basal Medium 2 | +2ng/ml FGF7<br>+0.25mM Vit. C<br>+100ng/ml EGF<br>+0.25uM Sant1<br>+0.1uM RA<br>+200nM LDN<br>+100nM TPB<br>+10ng/ml AA<br>+10mM Nicot.<br>+5uM RI |
| D14-D15 | Suspension |  | +0.25mM Vit. C<br>+100ng/ml EGF<br>+0.1uM RA<br>+100nM TPB<br>+25ng/ml FGF7<br>+100nM GSiXX<br>+10mM Nicot. |
| D16-D18 | Suspension |  | +25ng/ml FGF7<br>+0.25mM Vit. C<br>+100ng/ml EGF<br>+1uM RA<br>+100nM TPB<br>+100nM GSiXX<br>+10mM Nicot.<br>+100ng/ml FGF10<br>+10 ng/ul AA<br>+100nM LDN |

Here we describe a step-by-step protocol for the generation and culture of the pancreatic organoids derived from pancreatic progenitors cultured in 6-well plates and collected on Day 11 of the differentiation protocol. The general steps of cell dissociation, aggregation and suspension culture are similar to those described in Barsby et al. However, the timing and culture media have been modified and optimised to induce exocrine differentiation and branching morphogenesis instead of endocrine differentiation. The reagents required for these steps are listed in Table 2.

### Day 11

#### Preparation of the Aggrewell microwell system (24-well plate)

- Add 500μL of anti-adherence rinsing solution in each well and centrifuge at 1300g for 5min.
- Remove the solution by aspirating the anti-adherence solution and add 500μL of MCDB131 medium.

#### Dissociation and collection of pancreatic progenitors

- Wash pancreatic progenitors (cultured in a 6-well plate) by adding 1ml of Versene to each well and removing it after 1min.
- Add 1ml of TrypLE to each well and incubate for 8min at 37°C.
- Keep at 37°C until mild-tapping of the plate causes cell detachment.
- Add 1ml of MCDB131 medium per well and gently flush until most cells are detached.
- Add another 1ml of MCDB131 medium per well and collect cells from all wells into a 50ml tube.
- Count cells
- Centrifuge the 50ml tube with the cell suspension at 300g for 3min.
- Aspirate supernatant and gently resuspend cells in pre-warmed Basal Medium 2 supplemented with the cytokines and small compounds listed in Table 3 (D11-D13).
- Aspirate MCDB131 medium from the Aggrewell plate, replace with 2mL D11-D13 medium, and seed pancreatic progenitors at a density of 1200 cells/microwell.
- Centrifuge plate at 100g for 3min.
- Incubate plates in a humidified cell culture incubator (37°C, 5% CO_2_).

### Day 12-13

#### Media changes

- Carefully remove 1ml of medium from each well by keeping the plate horizontal and aspirating slowly with a pipette tip while taking care not to dislodge the clusters at the base of the well.
- Add 1ml of fresh pre-warmed D11-D13 medium very gently on the top side of the wells.

### Day 14

#### Prepare D14 culture medium

*On this day, cell clusters collected from 2 wells of the Aggrewell plate are pooled in one ultra-low attachment (ULA) well of a 6-well plate*.

-Prepare enough D14-D15 medium (Basal Medium 2 with supplements listed in Table3) to add 5ml per well of the ULA 6-well plate.

#### Transfer of pancreatic progenitor clusters to suspension culture

- For each well of the Aggrewell plate, carefully remove 1ml of medium and gently flush it out on the edge of the well to dislodge cell clusters from the microwells.
- Coat the pipette tip used to transfer the clusters with medium.
- Swirl the Aggrewell plate to pool the cell clusters toward the centre of the well and transfer them into a ULA plate. Pool cell clusters collected from 2 wells of the Aggrewell plate into one ULA well.
- Once all cell clusters have been transferred into the ultra-low attachment plate, swirl this plate gently in one direction to pool the clusters into the centre.
- Remove most of the medium by making sure not to aspirate cell clusters, which should be visible by eye.
- Add 5ml of D14-D15 medium per well.
- Place the plate in a humidified cell incubator (37°C, 5% CO_2_) on a rotating platform (100RPM).

### Day 15

- Swirl the ULA plate to centre the cell clusters in the wells.
- Remove 4ml of medium by aspirating from the well edge and replace with fresh D14-D15 medium.
- Place the plate back into a humidified cell incubator (37°C, 5% CO_2_) on a rotating platform (100RPM).

### Day 16-18

- Swirl the ULA plate to centre the cell clusters in the wells.
- Check the formation of branches as a marker of efficient organoid maturation.
- Remove 4ml of medium by aspirating from the well edge and replace with fresh D16-D18 medium.
- Place the plate back into a humidified cell incubator (37°C, 5% CO_2_) on a rotating platform (100RPM).

### Day 19

Proceed with experiments using the pancreatic organoids or follow the steps below to fix the organoids.

- Swirl the ULA plate to centre the organoids in the wells.
- Collect and pool organoids in a tube.
- Centrifuge at 200g for 2 min.
- Wash in PBS and centrifuge again before fixing in 4 % paraformaldehyde for 30 minutes at 4°C.
- Keep in PBS at 4°C.

### Immunofluorescence and RT-qPCR

#### Immunostaining of cells

In addition to the 6-well plates that were used to generate pancreatic progenitor-derived organoids, hiPSCs were also cultured in 24-well plates. These plates were used to confirm the efficiency of hiPSC specification into pancreatic progenitors by immunostaining of acinar and ductal progenitor markers at D14. Cells were washed in PBS, fixed for 20 minutes at 4°C with 4 % paraformaldehyde and washed again. Fixed cells were then blocked for 45min in a blocking buffer made of PBS, Donkey serum (3%) and Triton-X (0.1%). Primary antibodies were added into the blocking buffer at dilutions listed in Table 4 and incubated overnight at 4°C. The next day, cells were washed multiple times in PBS with Triton-X (0.1%) and incubated for 1 hour in the dark with the secondary antibodies diluted in blocking buffer (Table 4) and counterstained with Hoechst. Before imaging, cells were washed multiple times with PBS.

**Table 4.** List of reagents to perform immunostainings of cells and organoids.

| Reagent | Source |
| --- | --- |
| Guinea pig anti-PDX1 (1:300) | Abcam (Cat # AB47308) |
| Mouse anti-NKX6.1 (1:400) | DSHB (Cat #F55A10) |
| Rabbit anti-AMYLASE (1:400) | Merck (Cat #A8273) |
| Hoechst 33342 (250 ng/mL) | Invitrogen (Cat #H1399) |

#### Immunostaining of organoids

Throughout the staining procedure, solutions were changed by centrifuging organoids at 200g for 2min, removing the supernatant and adding the next solution.

First, fixed organoids were blocked for 24h in a blocking buffer made from PBS, Donkey serum (3%) and Triton-X (0.5%) at 4°C on a rotating platform. Primary antibodies were added to the blocking buffer at dilutions listed in Table 4 and left to incubate for 24h in similar conditions. The next day, organoids were washed three times for 45min in PBS with Triton-X (0.1%), and then incubated for 24h in the dark at 4°C on a rotating platform with the secondary antibodies diluted in blocking buffer (Table 4) and counterstained with Hoechst. Before imaging, organoids were washed twice in PBS with Triton-X (0.1%) for 45min, and once in PBS.

#### Imaging and Image analysis

Immunostained organoids were placed in a 96-well plate with a glass bottom and imaged using a Zeiss LSM 700 confocal microscope. Cell numbers and immunostaining intensities were quantified using ImageJ^21^. The PDX1 intensity was quantified by measuring the mean pixel intensity in regions of interest (ROI) delineating the AMY+ and NKX6.1+ cells.

#### RT-qPCR

Cells from planar culture were dissociated and harvested with TrypLE and organoids/cell clusters were collected with a trimmed pipette tip pre-coated with 1mL medium previously removed from the well. These were centrifuged for 3 minutes at 300 RCF or for 2 minutes at 200 RCF, respectively. The supernatant was aspirated, and the pellets were washed once with PBS (and then frozen at -80°C). A High Pure RNA Isolation Kit (Roche) was used for RNA extraction, and all samples were treated with DNase to remove any genomic DNA. cDNA was synthesised using a Transcriptor First Strand cDNA Synthesis Kit (Roche). RT-qPCR was performed on a Lightcycle (Roche) with the FastStart Essential SYBR Green Master Mix (Roche). The 2-ΔΔCt method^22^ was used to evaluated relative gene expression and the housekeeping gene glyceraldehyde-3-phosphate dehydrogenase (GAPDH) was used for normalisation. cDNA dilution was modified in order to return Cts of approximately 20 for GAPDH, and missing Ct values were set to 40 when the threshold for detectable fluorescence was not met. Primers used for RT-qPCR are listed in Table 5.

**Table 5.**
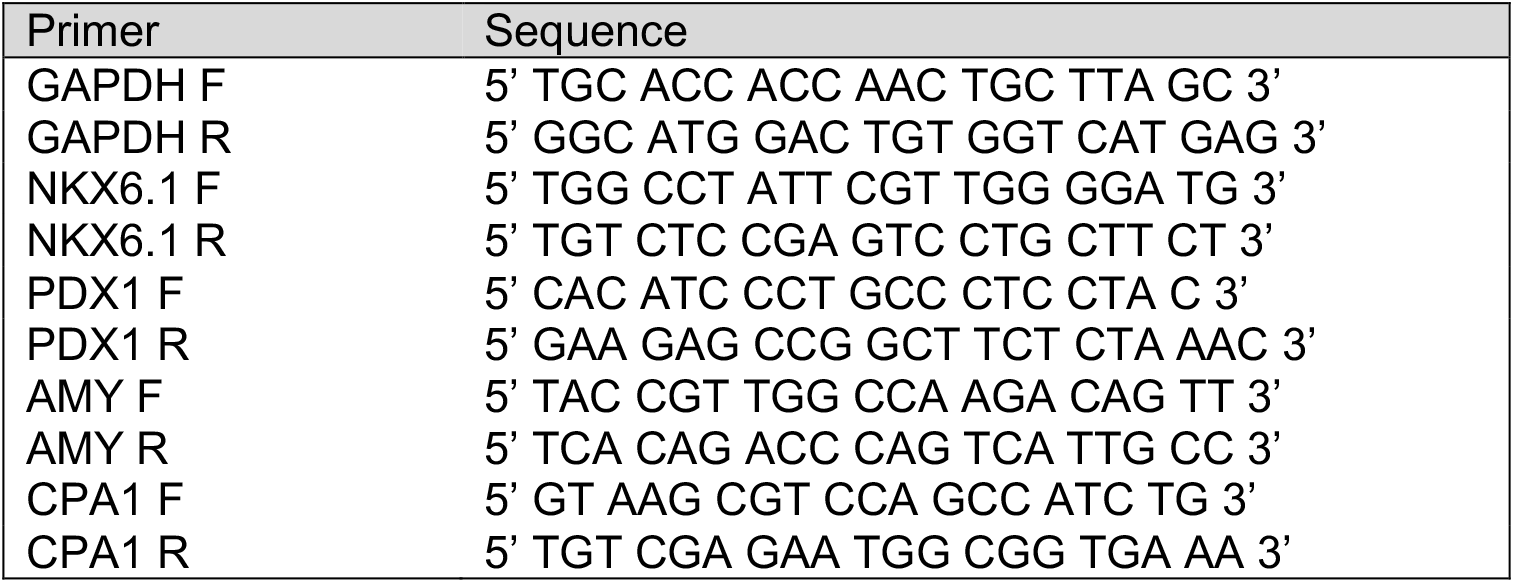
List of primers used to perform RT-qPCR on hiPSC-derived cells.

#### Quantification and Statistical Analysis

All statistical analyses were done using the R package rstatix. The elements shown in the boxplots are the 25% (Q1, upper box boundary), 50% (median, black line within the boxes) and 75% (Q3, lower box boundary) quartiles, and the whiskers represent a maximum of 1.5X the interquartile range. P-values were calculated using two-tailed t-tests. Normality was assessed using the Shapiro-Wilk normality test. Statistical significance was calculated using unpaired Student’s t-tests for data with normal distributions and Mann-Whitney U-tests otherwise.

## Results

### Generation of pancreatic progenitors

hiPSCs were differentiated into pancreatic progenitors as described by Barsby et al.^20^ with minor modifications, which are outlined in the materials and methods (Fig1A). The efficiency of pancreatic progenitor differentiation was monitored by assessing the morphology of differentiating cells with a phase contrast light microscope between D8 and D10 of differentiation and by performing immunostainings for the pancreatic progenitor markers PDX1 and NKX6.1 at D11 of differentiation (Fig1B). Pancreatic progenitors were predominantly double positive for PDX1 and NKX6.1 and clusters were efficiently formed within 24 hours in the Aggrewell plate.

**Fig 1.**
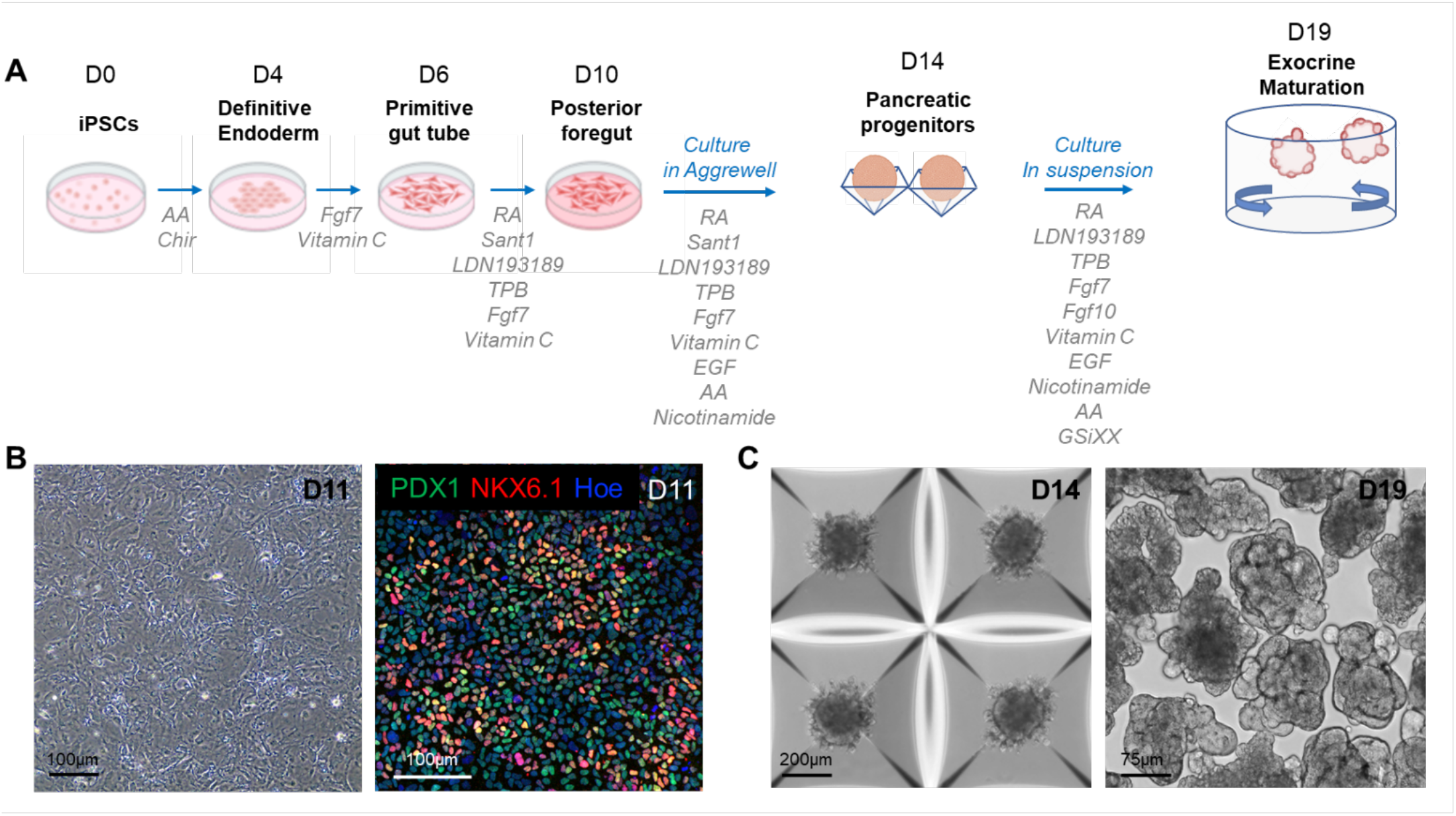
Differentiation of hiPSCs into branched pancreatic organoids. **A**. Schematic overview and media composition for the differentiation of hiPSCs into pancreatic organoids. The differentiation was carried out as a monolayer until day 11 (D11), then transferred to the Aggrewell microwell system from D12 to D14, and finally performed in suspension until D19. **B**. Representative phase contrast image (left panel) and confocal image (right panel) of D11 pancreatic progenitors immunostained with the PDX1 and NKX6.1 pancreatic markers. **C**. Representative phase contrast images of pancreatic clusters cultured in the Aggrewell microwell system on D14 of the differentiation (left panel) and in suspension on D19 of the differentiation (right panel). AA: Activin A, RA: Retinoic Acid.

### Generation of branched organoids

Dissociated pancreatic progenitors were left to aggregate in Aggrewell plates (Fig1C). After progenitor clusters formed, they were transferred to suspension plates for the final stage of differentiation using an optimized differentiation medium (see composition in Table 3). Specifically, we supplemented the medium with GSiXX, a Notch inhibitor known to promote pancreatic cell differentiation^38^, along with FGF7 and FGF10, which stimulate growth and morphogenesis of the exocrine pancreas^35^. In contrast to Barsby et al., we excluded factors known to induce endocrine differentiation, such as thyroid hormone receptor agonists, TGF-β antagonists and Betacellulin. This medium, combined with the suspension culture format, triggered primary branch formation in the organoids between D17 and D18. 93.5% (SD = 2.3) of organoids acquired a branched architecture by D19 of the differentiation, highlighting a high efficiency of morphogenesis.

To investigate the efficiency of our culture medium to induce exocrine differentiation in the organoids we performed immunostainings for tip and trunk markers (Fig 2A). Our wholemount immunostainings highlight that the stalk region of the organoid branches is positively marked by the NKX6.1 trunk marker, whereas the tip regions are marked by the Amylase acinar marker. Hence, the pattern of expression of both these markers are mutually exclusive and clearly define tip and trunk domains in the organoids. In addition, our quantifications show that the acinar domain strongly increases in the last stage of the differentiation-between D14 and D19-demonstrating that our protocol supports acinar cell differentiation (Fig 2B). Altogether, our differentiation protocol recapitulates the patterning of the pancreatic branches observed in vivo with acinar cells positioned at the tip of branches.

**Fig 2.**
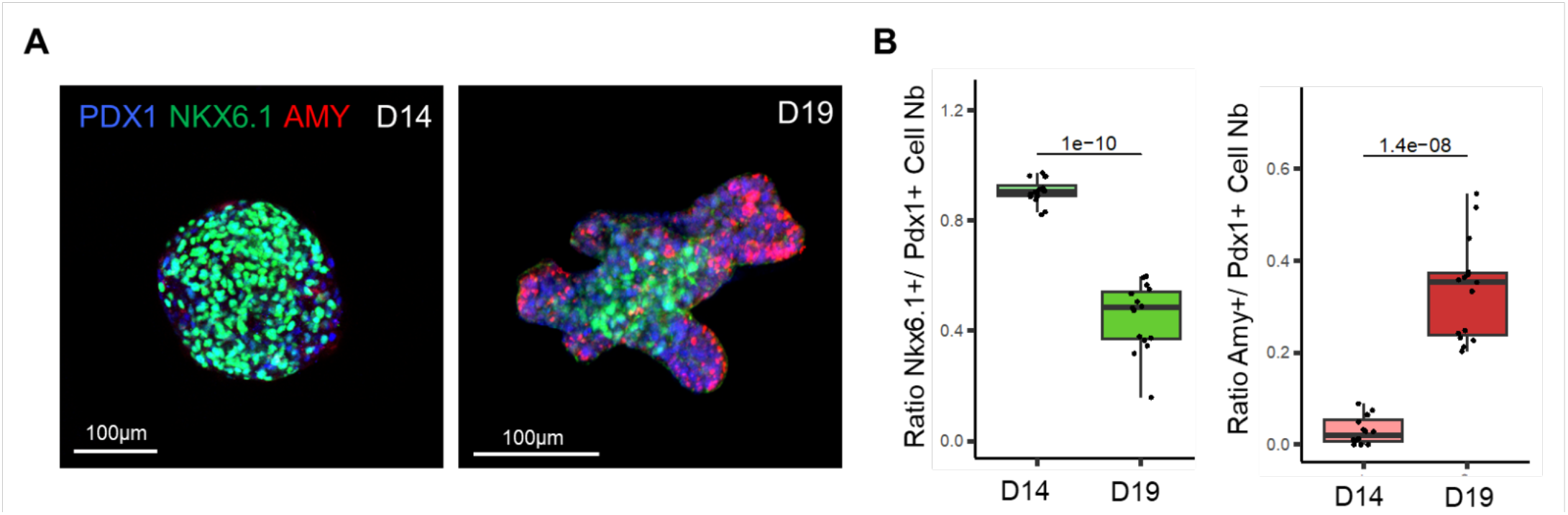
Physiological patterning of the pancreatic organoid branches. **A**. Representative confocal images of D14 pancreatic clusters (left panel) and D19 pancreatic organoids (right panel) immunostained for PDX1, NKX6.1 and Amylase (AMY), which respectively mark all pancreatic progenitor cells, trunk progenitor cells and acinar cells. **B**. Ratio of PDX1+ cells positive for the trunk marker NKX6.1 at D14 and D19 (left panel). Ratio of PDX1+ cells positive for acinar marker Amy at D14 and D19 (right panel). N=13-15 organoids per stage analysed, from N=3 differentiation experiments.

### Analysis of acinar and ductal cell differentiation

It was previously shown that during in vivo pancreas development PDX1 expression levels decrease in differentiating acinar cells^23^. To confirm that this process is recapitulated in our organoid system, we quantified the PDX1 staining intensity in AMY+ acinar cells compared to NKX6.1+ cells in organoids collected at D19 (Fig 3A). Our results show that PDX1 staining intensity is significantly lower in AMY+ acinar cells compared to NKX6.1+ cells at this stage, thus recapitulating the PDX1 expression decrease observed in vivo.

**Fig 3.**
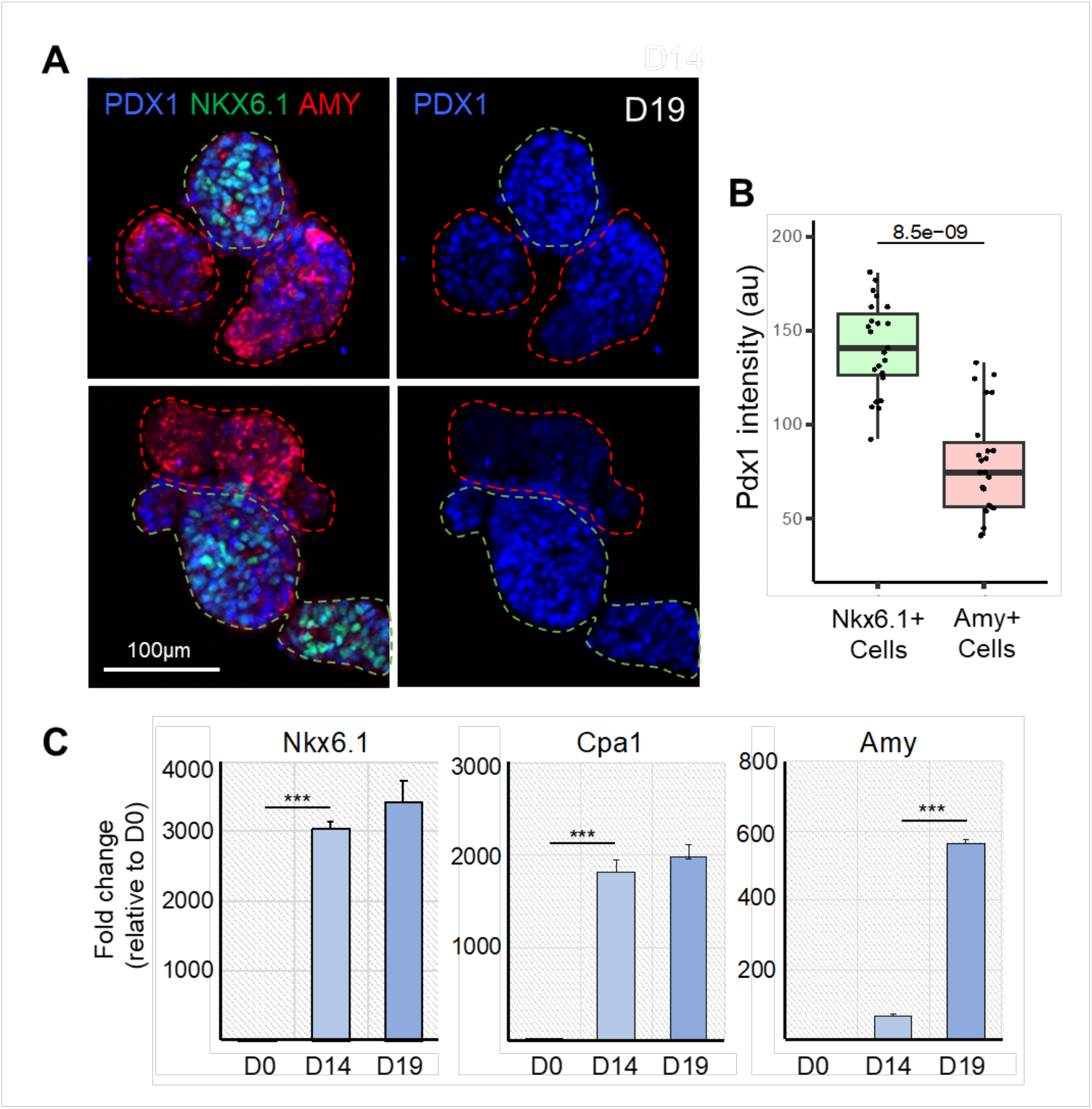
Differentiation of acinar cells in branched pancreatic organoids. **A**. Representative confocal images of D19 pancreatic organoids immunostained for PDX1, NKX6.1 and AMY, which respectively mark all pancreatic progenitor cells, trunk bipotent cells and acinar cells (left panel). PDX1 is shown alone in the right panel. Dotted green and red lines respectively delineate the NKX6.1+ trunk and AMY+ tip regions. **B**. Quantification of PDX1 staining intensity in the NKX6.1+ and AMY+ cells. N=17-22 organoids analysed, from N=3 differentiation experiments. **C**. Representative qRT-PCR analysis for the indicated pancreatic genes performed on cells, clusters and organoids collected at D0, D14 and D19 of the differentiation protocol. Data are represented as relative fold change. Values shown are mean ± SEM. *p < 0.05, **p < 0.01, ***p < 0.001; two-tailed paired t tests.

In addition, we performed RT-qPCR on dissociated cells collected at D0, D14 and D19 to investigate the efficiency of our protocol to induce cell differentiation, specifically the acinar fate. Our results highlight that differentiating cells acquire a pancreatic fate by D14, as evidenced by the strong expression of the pancreatic progenitor markers *NKX6*.*1* and *CPA1* (Fig 3B). Between D14 and D19, we observed that the level of *Amylase* expression strongly increased in organoids and *CPA1* expression is also maintained, indicating that our organoid culture medium promotes the differentiation of acinar cells. Altogether, the differentiation of acinar cells and the formation of a branched architecture demonstrates that our protocol enables the formation of pancreatic organoids with an exocrine identity.

## Conclusions/Discussion

The derivation of pancreatic organoids from either stem cells or primary tissue allows the possibility of replicating native pancreas organisation, morphology, and the emergence of distinct cell populations in health and disease^24^. Cell sources of human pancreatic organoids include hPSCs, primary cells from adult tissue, and foetal tissue obtained between 7-11 wpc^13^. Human PSC-derived pancreatic organoids are capable of giving rise to ductal or acinar cells that usually self-organise as polarised spheres^25,26^. However, the generation of human stem cell-derived pancreatic organoids with proper branched architecture and correct patterning of acinar/tip and trunk cell domains has been an ongoing challenge.

The advent of pancreatic organoids for use as an in vitro model follows many years of optimisation of in vitro pancreatic cell differentiation protocols^27–29^, which aim to mimic pancreas development. Most recently, a highly optimised protocol was developed^20^ that incorporated several existing protocols and made specific use of SHH and BMP inhibition, FGF and RA signalling activation, and the discovery that joint administration of Nicotinamide and EGF can increase pancreatic progenitor cell yield^30^. Nicotinamide has been shown to increase the proportion of PDX1+NKX6.1+ by inhibiting CK1 and ROCK^31^, while epidermal growth factor (EGF) is known to activate the MAP kinase pathway and is required for pancreatic progenitor proliferation^32^. This combination resulted in a heavily improved yield of PDX1+NKX6.1+ pancreatic progenitors relative to established protocols^20,33^. Incorporation of this protocol was essential for our work as it has also been reported that a differentiation efficiency resulting in fewer than 60% PDX1+NKX6.1+ pancreatic progenitors typically do not form aggregates of sufficient size when using the Aggrewell platform. This result also highlights the importance of adequately replicating in vivo complexity to achieve effective production of pancreatic cell types with high fidelity.

Here, we present a protocol for the generation of hiPSC-derived pancreatic organoids that bear a resemblance to the neonatal pancreas. We have shown that these organoids harbour regionalised tip and trunk domains. Prominent branching and organoid expansion were observed during the final stage of differentiation that may have resulted from supplementation with Fgf7 and Fgf10, which are known to be important for the growth and morphogenesis of the exocrine pancreas^34,35^. Fgf10 is also known to interact with and activate Notch to maintain pancreatic progenitor cell self-renewal^35–37^. However, Notch suppression is later required for proper lineage diversification and specification^38,39^, and several protocols use media supplementation with Notch inhibitor for the differentiation of PSCs into different pancreatic cell types in vitro^28,40,41^. Since in this work Notch inhibitor is applied from D14 onwards, our findings suggest that regionalisation is at least in part achieved by D14, after which Notch inhibition enables regionalised cell differentiation. The use of the Aggrewell microwell platform was a necessary intermediary step in creating an environment that facilitates cell interactions in 3D, without which branched organoid structures could not be generated. Moreover, our protocol eliminates the need for matrices such as Matrigel or collagen to culture pancreatic organoids. This is a significant advantage as it reduces the use of animal-derived reagents and minimizes the unpredictability of organoid differentiation efficiency due to reagent batch-to-batch variability.

Further optimisation will be able to determine whether the maturation of these organoids can be improved when cultured for extended periods of time. The fact that pancreatic cells dedifferentiate into embryonic-like states during pancreatic inflammation and PDAC^15^, which are the states replicated in our system, suggests that our organoid platform could allow *in vitro* modelling of many aspects of the diseases and enable preclinical investigations in a high-throughput setup while partially replacing animal models. Pancreatic organoids derived from pluripotent human stem-cells have indeed been used to study PDAC initiation and progression in culture by induction of the KRAS and GNAS oncogenes^26,42^. So far, these studies were performed on cystic organoids solely composed of ductal or acinar cells. Inducing the expression of the KRAS and GNAS oncogenes in our organoid platform would enable the research community to study the effects of these mutations in a system that both recapitulates the branched architecture of the pancreas and the patterning of its acinar and ductal compartments. Moreover, the high-throughput nature of this platform could allow the screening of drugs that reduce tissue alterations caused by oncogene induction and thus identify potential therapeutic targets for pancreatitis and PDAC.

## Acknowledgements

This work was supported by a fellowship from the NC3Rs (grant reference NC/V002260/1) to Jean-Francois Darrigrand. Abigail Isaacson was funded by the Wellcome Trust PhD programme ‘‘Cell therapies and regenerative medicine’’ (grant #108874/Z/15/Z).

Figure 1 was created with BioRender.com.

